# Modulation of the internal dynamics of the Homer1 EVH1 domain by putative autism-associated mutations

**DOI:** 10.64898/2025.12.17.694870

**Authors:** Fanni Farkas, Brigitta Maruzs, Zsófia E. Kálmán, Tomas Klumpler, Gyula Batta, Bálint Ferenc Péterfia, Zoltán Gáspári

**Author notes:** To whom Correspondence should be addressed: Zoltán Gáspári, Bálint Péterfia.

## Abstract

The EVH1 domain of the Homer1 scaffold protein interacts with the proline-rich region of Shank3, forming a key network within the postsynaptic density. Two mutations in EVH1, M65I and S97L, were suggested to be associated with autism spectrum disorder. Combining experimental and computational investigations, we show here that neither the overall structure nor the partner binding properties of the mutants exhibit substantial changes relative to the wild type. Compared to the S97L variant, the M65I mutant exhibits larger chemical shift perturbations both upon the mutation itself and during partner binding, and also shows signs of thermal destabilization. Integration of computational and NMR investigations suggests that both mutations perturb the μs-ms time scale internal motions of the EVH1 domain.

## Introduction

Homer1 is the most studied member of the Homer family. It is a scaffold protein in the postsynaptic density (PSD), found beneath the postsynaptic membrane of central excitatory glutamatergic synapses [1]. The PSD is a complex structure with high concentration of glutamate receptors, cell adhesion molecules, signaling enzymes and cytoskeletal elements [2]. The PSD is a dynamically changing structure whose composition is different during response to synaptic activity, development, learning and memory formation [3]. Pathological synapse development can contribute to different disorders (schizophrenia, mental retardation, Alzheimer’s disease and autism spectrum disorder) [3].

There are three mammalian genes encoding Homer1, 2 and Homer3. The cortical and CA1 hippocampal regions express mostly Homer1 which has 3 splice variants (Homer1a, -1b, -1c). Homer1c is also (named PSD-Zip45/Vesl) found in the PSD [4]. Homer1c in PSD plays a crucial role in receptor clustering and trafficking, intracellular complex formation, and calcium homeostasis [1]. It interacts with proline-rich motifs in various proteins such as mGluRs, actin, Oskar, RYRI, IP3Rs, Drebrin and Shank/ProSAP/SAPAP [4] [1] [5] [6]. This interaction is mediated by its N-terminal globular EVH1 (Enabled/vasodilator-stimulated phosphoprotein (Ena/VASP) homology 1) domain. Besides EVH1, it has a C-terminal coiled-coil segment and a 77-residue long disordered region between them. EVH1 has 7 beta strands and a C-terminal alpha helix. EVH1 forms an interaction with consensus sequences FPxoP and PPxxF [7] [5]. Homer1c binds its ligands by Gly89, Val85, Gln76, Phe74, The70 and Trp24 amino acids.

Our previously determined solution structure of the wild-type EVH1 domain (PDB ID: 9QUX) exhibits high overall similarity to the other available EVH1 structures and shows excellent agreement with SAXS measurements [8]. However, it exhibits subtle conformational differences relative to X-ray structures in several loop regions, including regions 17–29 and 40–43, involved in partner binding. In addition, in the NMR structure, the indole ring of Trp24 adopts a nearly perpendicular orientation relative to the Homer1 EVH1 crystal structures (1DDV and 1DDW [5], suggesting a possible reorganization of the key aromatic cluster (Phe14, Trp24, Phe74) during partner binding. These findings highlight the structural plasticity of the binding site and its potential role in accommodating diverse pro-rich ligands [8].

Shank is one of the interaction partners of Homer1c. It is a multidomain synaptic scaffold protein [9]. Starting from the N terminal depending on the alternative splicing, Shank proteins possess an ankyrin repeat region, an SH3 domain, a PDZ domain which is the main interaction partner in Shanks and participates in the NMDA-PSD-95-GKAP-Shank complex. Besides, Shank has a long disordered proline-rich domain that interacts with the EVH1 domain of the Homer protein. Lastly it has a C-terminal SAM domain.

The Homer EVH1 domain recognizes PPxxF motifs in its interaction partners (metabotropic glutamate receptor (mGluR1 and mGluR5), IP3 receptors (IP3R), Shank, transient receptor potential canonical (TRPC) family channels, drebrin, oligophrenin, dynamin3, CENTG1, and ryanodine receptor). The consensus PPxxF sequence is different in the known Shank paralogs. In Shank1 its sequence is PLPPPLEFSN, in Shank2, there are two putative sites: PLPPPLEFAN and FLPPPESFDA, and in Shank3, the sequence is LVPPPEEFAN [10]. Through Shank, Homer crosslinks with the mGluR as well. Homer regulates the postsynaptic localization and intracellular signaling of mGluR1 alpha/5and also modulates its traffic from the cytoplasm to the plasma membrane.

The SHANK3 gene was identified as an autism spectrum disorder (ASD)-risk gene and recently was found that it is one of the most frequent causes of ASD [9].

Autism spectrum disorder is a neurodevelopmental disability condition in early childhood affecting 1% of the children [11] [12]. It is associated with repetitive behaviour, impaired communication and social interaction, and a limited range of interests [12] [13]. In relation to ASD copy number variants (CNVs) and mutations in individual genes have been documented [12]. Homer1 has been linked to multiple neurological disorders such as Alzheimer’s disease schizophrenia including ASD [1] [14]. The HOMER1 gene was identified as a novel autism-risk gene. Homer1 participates in the downstream mGluR signaling, alteration in this pathway contributes to non-syndromic autism [12]. In the HOMER1 gene five missense point mutations (M65I, S97L, P142L, R323H, 3’UTR) were reported, based on the sequence, M65I and S97L (DNA: 195G>T resulted M65I and DNA: 290C>T resulted S97L) found in the EVH1 domain of Homer1 [12]. To our knowledge, the effect of the mutations on the structure and function of the EVH1 domain has not yet been investigated.

In our previous study, complete backbone and partial sidechain NMR chemical shift assignments were obtained for the wild type EVH1_1-118_ domain. In this study, to explore the effect of putative ASD-related mutations, we report the expression, purification and resonance assignment of two murine Homer1 EVH1 mutant (M65I and S97L) domain constructs. We present the analysis of the backbone dynamics and the partner binding properties of these variants and compare these to each other and the wild type EVh1 domain. Using NMR and CD spectroscopy, SAXS, thermal shift assay, biolayer interferometry and molecular modeling, our integrated analysis suggests that the mutations primarily affect the characteristic internal dynamics of the domains rather than their overall structure or ligand binding at the canonical site.

## Materials and Methods

### Cloning, protein expression and purification

After successful production and purification of the wild-type (WT) Homer1 EVH1 protein, using the segment 1-118 from *Mus musculus* (UniProt ID: Q9Z2Y3) (see details in our previous article: [8], two autism-associated point mutations, M65I and S97L, were introduced in its sequence with *in vitro* PCR mutagenesis. Proteins were expressed in unlabeled and ^15^N and ^15^N, ^13^C labelled media as well. For unlabelled, LB medium was used. For isotopically labelled protein freshly prepared M9 medium was used [15] supplemented with 0.2x Trace element mix [16], 2.5 g/L ^15^NH_4_Cl (Cambridge Isotope Laboratories, Cambridge, MA), and 4 g/L unlabeled glucose or [^13^C]-D-glucose in the case of double labeled version. With ultrasonic homogenization (30 min, 30% pulse, 30% power) the bacteria cell wall was destroyed. To make protein of interest available, the following purification steps were used; first IMAC was performed using 5 ml Bio-Scale™ Mini Nuvia™ IMAC Ni-affinity column (Bio-Rad). This column binds the His-tag in the target protein, therefore with a washing step other unbounded proteins can be rid of. To cleave the His-tag from the protein of interest, TEV protease was added. The second purification step was the Ion Exchange Chromatography by using the Bio-Scale™ Mini Macro-Prep® High S column. With a reverse IMAC step the cleaved His-tags were eliminated from the target protein. Sample was concentrated (∼9 mg/ml) and finally to get an NMR compatible pure sample without any additional contamination a third purification step was applied; Size Exclusion Chromatography by using the SEC70 analytical gel chromatographic column (Bio-Rad). The recombinant EVH1 proteins were eluted in the NMR compatible low salt NaPi buffer (50 mM NaPi; 20 mM NaCl, 0.02% NaN_3_; pH 7.4) and concentrated again for NMR measurement. To determine its purity and molecular weight, SDS-PAGE (Sodium-dodecyl-sulfate polyacrylamide gel electrophoresis), and NanoDrop2000 were used.

Molecular mass of the protein constructs was analyzed by HPLC-MS using a Shimadzu LC-MS-2020 device equipped with a Reprospher-100 C18 column and a positive–negative double ion source (DUIS±). A quadrupole MS analyzer in a range of m/z 50–1000 was used. For gradient elution, 0.1% formic acid in water (as eluent A) and 0.1% formic acid in acetonitrile (as eluent B) were used.

### Biolayer Interferometry (BLI)

Streptavidin biosensors (Sartorius) were hydrated in a kinetic buffer (50 mM NaPi pH 7.4, 20 mM NaCl, 1% BSA, 0.1% Tween 20) at 25°C for 10 min before the experiments. The concentration of Tween 20 and BSA in this buffer was increased to 0.1% and 1% respectively in order to minimize the strong aspecific binding of the mutant EVH1 constructs to the biosensor. Kinetic assays were performed using a BLItz system (ForteBio, USA). Biotinilyated Shank3 peptide ligand (BioBasic, Toronto, CA) with the sequence LVPPPEEFANG (corresponding to residues 1381-1391 of UniProt entry Q9BYB0 representing human Shank3 but also occurring with identical sequence in the mouse and rat Shank3 proteins) was anchored to the biosensor surface for 60 sec at 2 mM concentration. For reference, only the kinetics buffer was added for 30 sec. Five different EVH1 analyte concentrations (∼10-, 5-, 2,5-, 1.25- and 0 μM) were used for the 30 sec long association step of the runs. The equilibrium dissociation constant (Kd) was determined from the BLI data using the data analysis software’s global fitting method. For reference, the curve with 0 μM analyte was used.

### Thermal Shift Assay

The Thermal Shift Assay (TSA), also known as Differential Scanning Fluorimetry (DSF), was used by rPCR (Thermo Scientific, PikoReal2 qPCR) to determine thermal denaturation of the unbinded and peptide binded WT and mutant EVH1 domains. It detects fluorescence changes from a dye (SYPRO Protein Gel Stains, Thermo Scientific) that forms a binding with the hydrophobic region upon unfolding. It is a technique widely used to study protein stability and ligand binding because of its simplicity, low sample requirement, and high-throughput capability. Melting temperature (Tm) shifts indicates the stabilizing or destabilizing effect of a ligand or environmental condition on the protein [17]. For TSA measurement 0.375 mg/ml protein (WT, M65I, S97L), 186 mM Shank3 peptide (BioBasic, Toronto, CA) and 5x SYPRO were used. As a protocol setting temperature was increased from 25 Celsius degree to 90 with 12 sec hold time, temperature increment after hold 0.3 °C.

### ECD spectroscopy

Far-UV CD spectroscopy measurement was performed with JASCO J-1500 CD spectrometer (JASCO Corporation, Tokyo, Japan) in 195-250 nm spectral range, 50 nm/min scanning speed, 1 nm bandwidth, 0.2 nm step size, 0.5 s response time and 3 scans of accumulation and baseline correction. It was recorded in a 1 mm pathlength cuvette (J/21 quartz). Sample concentrations were 10 μM in 50 mM NaPi, 20 mM NaCl, pH 7.4 puffer. In order to check temperature dependence, the 20°C-60°C interval was used, measurements were done at ten degree steps.

### Small Angle X-ray scattering measurements

Samples containing the M65I and S97L mutant EVH1 domains were prepared as described above but without isotope labeling. The concentrations for the SAXS measurements were set to 4 and 2 mg/ml. SAXS measurements were performed on a Rigaku BioSAXS-2000 instrument at CEITEC (Brno, Czech Republic), equipped with a HyPix-3000 detector, at a sample-to-detector distance of 0.48 m. The scattered intensity was recorded over a q-range of 0.008–0.65 Å⁻¹, where q=4πsinθ/λ, 2θ is the scattering angle, and λ=1.54 Å. Six frames of 10 min each were collected at a sample temperature of 25 °C. The raw data were normalized to the intensity of the transmitted beam and radially averaged. Each individual frame was inspected for radiation damage prior to averaging. Finally, the scattering contribution of the solvent blank was subtracted to obtain the sample scattering profile. Structure-based calculation of SAXS curves was performed with Pepsi-SAXS [18] using the structural models built based on the wild-type structure after equilibration (see the molecular dynamics calculation section below) containing the expression tags (GSH) at their N-terminus to properly reflect the samples actually used for the measurements.

### Nuclear magnetic resonance measurements

All experiments were performed on a Bruker Avance NEO 700 MHz spectrometer at 298 K. For the resonance assignment of the M65I and S97L variants, 500 μl samples containing 50 mM sodium phosphate buffer, 20 mM NaCl, 0.02% NaN_3_ at pH 7.4 with 6% D_2_O were used with protein concentrations of 491 μM for the M65I and 560 μM for the S97L variants, respectively. ^15^N-^1^H HSQC, ^13^C-^1^H HSQC as well as standard triple-resonance spectra were recorded: HNCO, HN(CA)CO, HNCA, HN(CO)CA, HNCACB and HN(CO)CACB. HCCH-TOCSY, HCCCONH. NMR spectra were processed in Bruker TopSpin, and analysis was performed in CCPNMR v3.2.3 [19] and NMRFAM-Sparky 1.413 [20]. neighbor-dependent secondary chemical shifts were calculated by the method described by Tamiola et al [21] as included in the CoNSEnsX^+^ server [22].

For backbone ^15^N relaxation measurements, ^15^N labeled samples were used, containing 207 μM protein for the M65I and 228 μM for the S97L variant. Standard Bruker pulse sequences in pseudo3D implementation were used. The delay times were 40, 80, 120, 160, 200, 300, 400, 600, 1200 and 2400 ms for T_1_ and 1, 2, 3, 4, 5, 6, 7, 9,12, 15, 18 cycles of a 16.96 ms loop (from the parameters p30 and d21 in the Bruker pulse sequence) for T_2_ measurements. For these experiments, a delay time (D1) of 4s was used. ^1^H-^15^N heteronuclear NOE was measured in an interleaved fashion with a delay time of 8s. The obtained spectra were first processed with TopSpin. Peak intensity calculation and curve fitting was performed with NMRFAM-Sparky 1.413 [20]. For model-free analysis, the program Tensor2 was used [23]. Cross-peaks exhibiting substantial overlap were excluded from the estimation of the correlation time and model-free analysis. The rotational correlation time was estimated using data only for residues with T_1_/T_2_ values within 1 standard deviation of the average. The analysis was performed to ensure compatibility with our earlier investigation of the wild-type domain [8].

To explore possible phenomena associated with slow-timescale dynamics in more detail, ^15^N CEST measurements [24] were performed on ^15^N labeled samples for both mutants plus the wild-type EVH1 domain. The delay time was 1s and the irradiation time was set to 4s. Initial analysis of the data was conducted using ColmarVista [25]. As no clear effects were detected upon visual inspection of the peak intensities, no further analysis of these data were performed.

NMR titration of the ^15^N-labeled WT1-118 (571 μM), M65I1-118 (509 μM) and S97L1-118 (352 μM) Homer EVH1 was performed with a short Shank3 peptide (BioBasic, Toronto, CA) corresponding to the immediate binding motif (LVPPPEEFANG) (+ 10 μl / 4.5 mM peptide / titration point). Ten titration points were measured for each variant. Data were analyzed using CCPNMR v3.2.3 [19].

Amide NH chemical shift perturbations observed for the mutants and upon titration with the Shank3 peptide were calculated using the formula 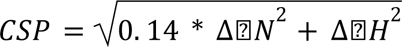.

### Computational structure modeling and analysis

For molecular dynamics calculations, the initial wild-type structure was derived from the first (representative) model of our previously determined Homer1 EVH1 NMR structure (PDB ID: 9QUX). Mutations were introduced with FoldX [26]. Molecular dynamics simulations were performed with GROMACS (version 2023.2) [27] using explicit solvent classical MD. Three parallel simulations for the wild-type and the two mutant domains were performed. AMBER99SB-ILDN [28] were used as a force field. During the simulation the box setting was a minimum distance of 1.0 nm from the box edge. Two rounds of equilibration were performed using 1 ns simulations with position restraints on the heavy atoms first using volume coupling (NVT ensemble) and then temperature coupling (NPT ensemble). From the 1 ms production run, 500 snapshots were taken. Computations were performed on the Komondor supercomputer. RMS fluctuations were calculated with GROMACS rmsf command. Secondary structure of the models was analyzed using DSSPcont [29]. Non-covalent interactions were analyzed using RING [30].

Gaussian network model analysis was performed using Prody [31] on a model of the wild-type protein from which the expression tag was removed and was renumbered accordingly. Structure visualization was done with ChimeraX [32]. Python3 was used for further data processing and Gnuplot and MSOffice Excel for data visualisation.

Human EVH1 sequences were downloaded from Interpro [33] and sequences aligned in Jalview [34] using ClustalO [35]. Orthologous sequences were downloaded from OMA [36].

## Results and discussion

### 1H-15N HSQC spectra indicate different effects of the mutations

Throughout this work, we use the residue numbering corresponding to the UniProt entry Q9Z2Y3. We note that relative to this, the numbering of our previously deposited Homer1 EVH1 NMR structure (PDB ID: 9QUX) is shifted by 3 residues due to the presence of the remnants of the expression tag.

The S97L and M65I variants of the Homer1 EVH1 domain could be successfully expressed using the protocol applied for the wild-type domain. Polyacrylamide gel electrophoresis and mass spectrometry measurements indicate that the obtained samples conform to the expected molecular mass of the variants. NMR resonance assignment could be completed for both mutants with over 87% completion for backbone atoms. Chemical shifts along with ^15^N relaxation data have been deposited in the BMRB with IDs 53479 (M65I) and 53480 (S97L). In general, most of the amide peaks of the S97L variant align well with those of the wild type EVH1 domain, while for the the M65I variant, the ^1^H-^15^N HSQC spectrum indicates the alteration of the chemical environment for the majority of the residues (Figure 2A). For both mutants, amide cross-peaks of the residues in and near the mutated positions show marked alterations. In the case of the M65I mutant, the affected region is longer, with some residue positions in the segment 60-78 exhibiting larger chemical shift perturbations. This observation suggests that the effect of the two mutations is profoundly different and prompted further investigations on the structures of the two variants (Figure 2B, Figure S1).

**Figure 1.**
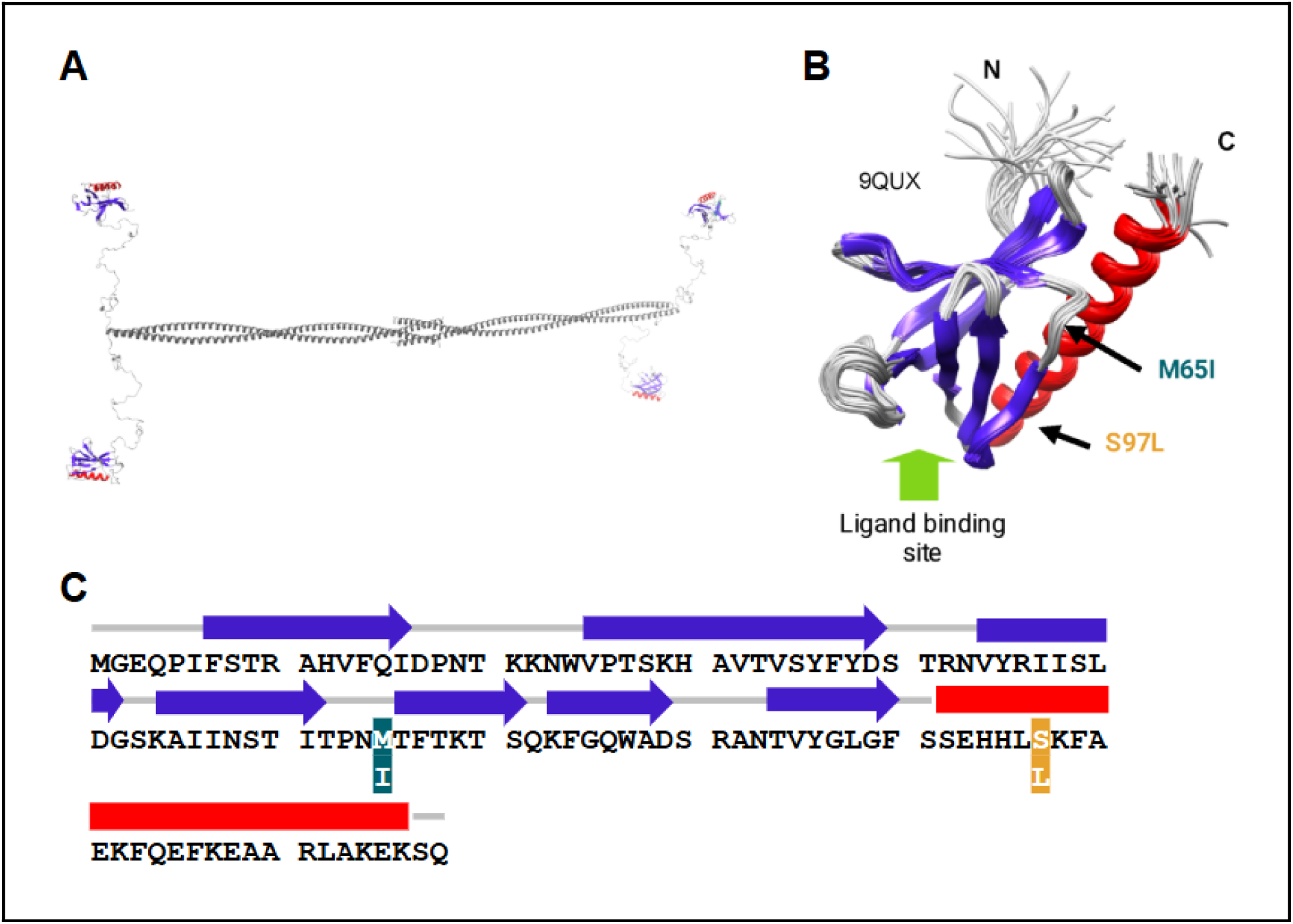
Overview of the structure of the Homer1 molecule and its EVH1 domain. **A)** Architecture of the full-length Homer1 tetramer (model from Kálmán et al. 2025, [8]). **B)** Solution structure of the Homer1 EVH1 domain (PDB ID: 9QUX). The ligand-binding site, as well as the positions of the M65I and S97L mutations are indicated. Figure prepared with ChimeraX. **C)** Sequence of the Homer1 EVH1 domain with the secondary structure elements and the mutations highlighted.

**Figure 2.**
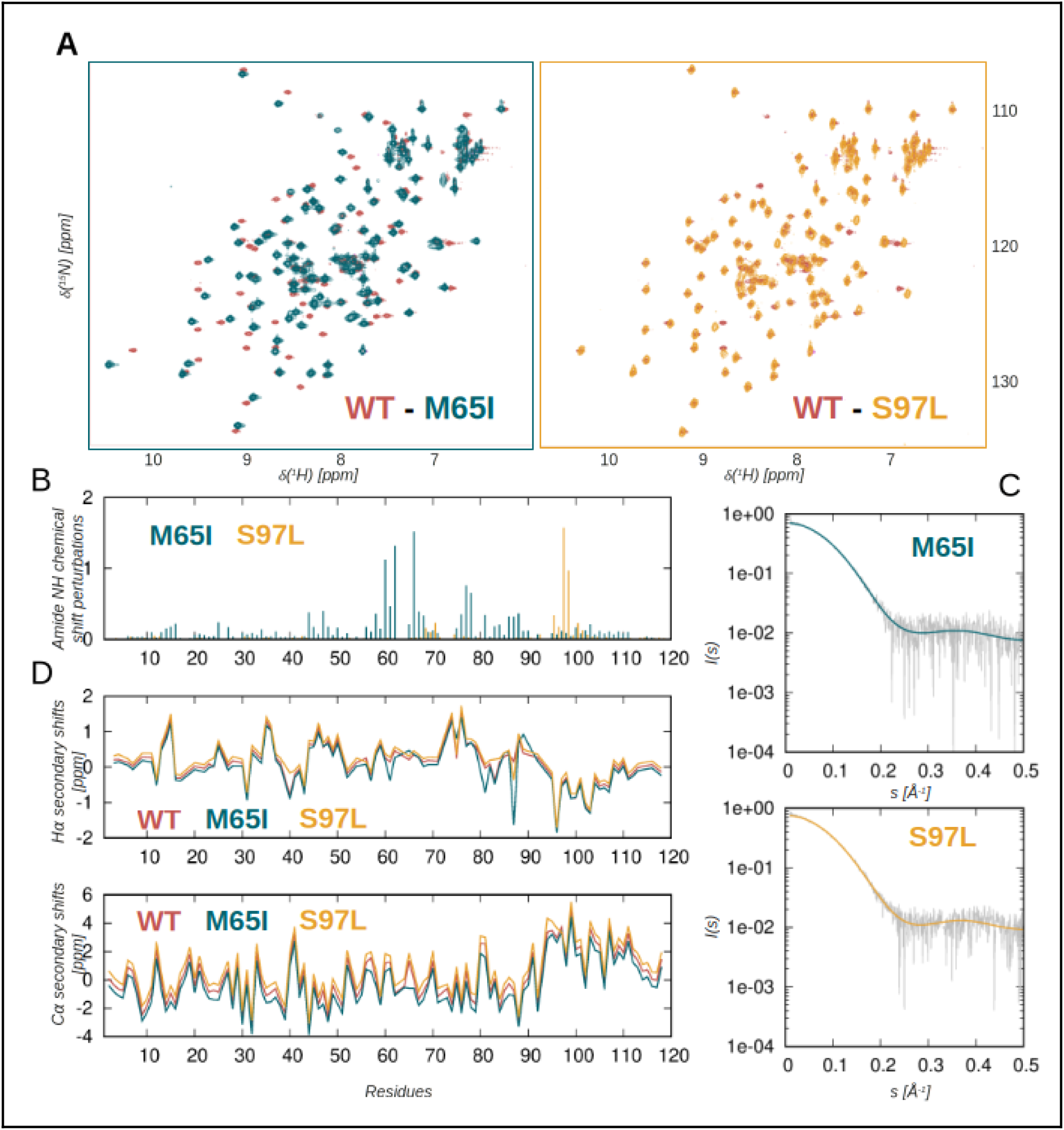
Structural analysis of the EVH1 domain variants investigated. **A**) Overlaid ^1^H-^15^N HSQC spectra of the mutant EVH1 domains with the wild type. Peaks are color-coded as indicated. **B**) Amide NH chemical shift perturbations (CSPs) for the two mutants along the sequence. CSPs were calculated as: 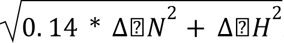. **C**) SAXS fits to structural models of the mutants built based on the wild type and having essentially the same overall structure. The experimental curves are shown in grey, those calculated for the structural models in color. **D**) Secondary Hα and Cα chemical shifts of the wild type and mutant variants. The interconnecting lines are only shown for visualization purposes to emphasise the similar trends in the data. Data for the mutants are slightly shifted upwards and downwards to reduce the overlap of the lines and aid better visualization. Data for the wild type are from [8] (BMRB entry 34990)

### The overall structure of the mutated EVH1 domains is largely unaltered

Both circular dichroism spectroscopy and small angle X-ray scattering experiments indicate that neither of the mutants exhibits a structure substantially different from that of the wild-type domain. This was confirmed by fitting the SAXS curves to the structural models built based on our previously determined EVH1 structure. The χ^2^ value of the fit for the M65I variant is 1.12, and 1.10 for the S97L mutant (Figure 2C). SAXS results have been deposited in the SASBDB database with IDs SASDYW3 (M65I) and SASDYX3 (S97L). Analysis of the secondary Cɑ and Hɑ chemical shifts also confirms that the secondary structure elements remain largely the same relative to the wild type domain (Figure 2D). These data strongly suggests that the observed chemical shift perturbations are not due to a substantial structural rearrangement upon mutation of the M65 residue. Therefore, we set out to explore the thermal stability and internal dynamics of the mutants in more detail.

### The M65I variant shows thermal destabilization and a markedly two-step unfolding kinetics

The stability of the EVH1 domain variants was investigated by thermal shift assay experiments. Interestingly, all three variants investigated - the wild type, S97L and M65I mutants - exhibit two transition temperatures, but the lower one of these is indicated only by a relatively small peak in the case of the wild-type and the S97L mutant (Figure 3A). Surprisingly, the S97L variant shows a slightly decreased first but an elevated second transition temperature compared with the wild type domain. In sharp contrast, the melting curves for the M65I variant exhibit a higher peak at the first and a smaller peak at the second transition that occurs at a lower temperature than for the wild type domain.

**Figure 3.**
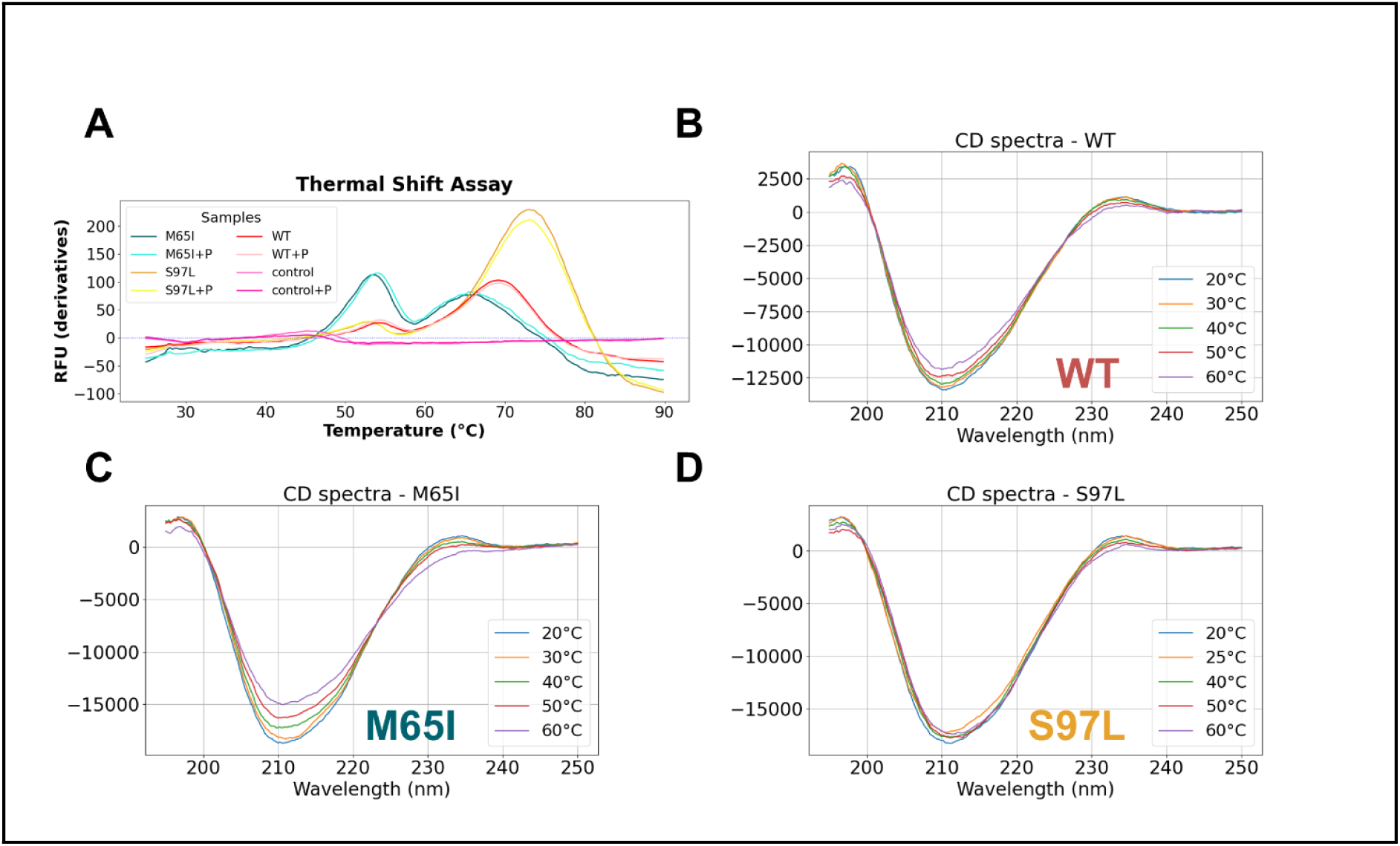
Temperature dependence of the structure of the EVH1 variants. **A)** Thermal shift assay curves for all three variants with and without ligands. Average curves from duplicate measurements are shown. **B)** wild type, **C)** M65I and **D)** S97L variants. The spectra for the wild type were taken from our previous work [8].

Intriguingly, binding to the Shank3 peptide causes only subtle alterations in the melting curves, indicating that the interaction does not cause any noticeable stabilization or destabilization.

Circular dichroism spectra do not indicate the presence of any substantial structural changes at higher temperatures, with the highest alterations observed again for the M65I variant (Figure 3B,C,D).

### NMR spectroscopy suggests different conformational response to ligand binding in the mutants

We tested the ability of the variants to bind a partner peptide derived from Shank3. Biolayer interferometry measurements indicated that the mutants bind the peptide with a comparable affinity to that of the wild type domain, with S97L showing a slightly stronger and M65I a slightly weaker binding (Table 1 and Figure 4A). Even so, the difference between the strongest binder, S97L, and the weakest, M65I, is still only approximately only twofold, indicating that the overall strength of the binding is not substantially affected by the mutations.

**Figure 4.**
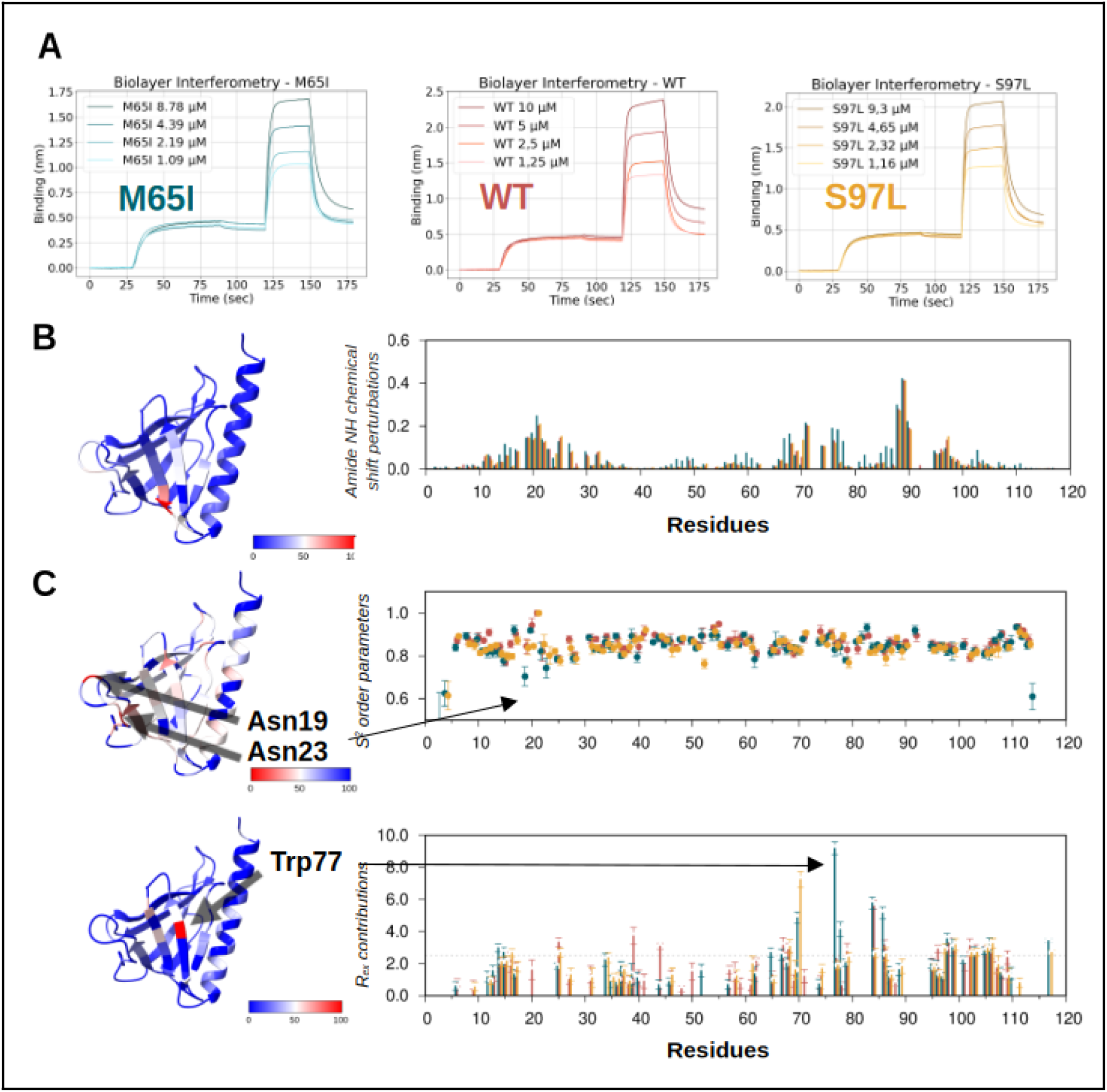
Ligand binding and internal dynamics of the EVH1 variants investigated. **A**) Biolayer interferometry assay probing the binding of the EVH1 domain variants to the Shank3 peptide. **B**) Amide NH chemical shift perturbations observed upon ligand binding for the variants. CSPs were calculated as: 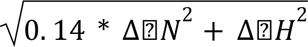. The structure on the right is colored according to the changes in the M65I variant.. **C**) S^2^ backbone order parameters and fitted R_ex_ values of the variants investigated. Structures on the right show residues with notable differences to the wild type in the M65I mutant. To highlight the lowest relevant S^2^ values on the structural model outside the termini, the color range was applied between 0.7-1.0 with lower values uniformly colored red. Data for the wild type are from [8] (BMRB entry 34990). Structure depictions prepared with ChimeraX.

**Table 1.** Binding affinity of the variants to the Shank3 peptide as determined by biolayer interferometry.

| Variant | WT | M65I | S97L |
| --- | --- | --- | --- |
| Global Kd [μM] | 5.15 | 8.88 | 3.88 |
| R<sup>2</sup> (coefficient of fit) | 0.99 | 0.99 | 0.98 |

We have also explored the response of the domain variants to peptide binding by NMR spectroscopy. As expected, a number of amide cross peaks corresponding to residues near the ligand binding site exhibit changes in their position in the ^1^H-^15^N HSQC spectra. Although the overall pattern is largely similar in all three variants (Figure 4B).

### 15N relaxation analysis suggests the presence of distinct slow-timescale motions in the variants

To explore the internal dynamics of the mutant EVH1 domains, ^15^N relaxation measurements were performed and evaluated with model-free analysis (Figures 4C and S2-S5). The results were compared with our previous analysis of the wild-type domain. Interestingly, residue Asn23 in the ligand binding site exhibits a heteronuclear NOE value above 1 in all three variants, as do some terminal residues. These were excluded from the final model-free analysis, but for Asn23, an internal residue, this phenomenon is worth noting as it might indicate complex dynamical behavior for this residue. Backbone S^2^ order parameters of the mutants align generally well (Figure 4C and S4) with those determined for the wild type domain. The obtained model-free fits were in some cases regarded as unreliable, either with very high (>1) S^2^ or S^2^f values and/or no robust fit to either of the motional models in Tensor2. However, most of the affected residues are in the terminal regions where increased mobility is expected. In the core of the domain, the largest difference relative to the wild type occurs in the segment between residues 19-15 where, in the case of the M65I mutant, two asparagine residues, Asn19 and Asn23, have lower S^2^ values than the wild type (Figure 4C and S4). Interestingly, in S97L, Lys21 is fitted with an S^2^ of 1. These residues are very close to the ligand binding site, and it is notable that the two mutants show opposite changes in their ps-ns time scale dynamics in this region. However, the largest difference in the internal dynamics of the variants is the different distribution of residues with a fitted R_ex_ contribution (Figures 4C and S5), also apparent in their R_2_ relaxation rates. Somewhat subjectively, we have defined a threshold of 2.5 Hz above which we consider the R_ex_ contributions substantial, but for most of the cases discussed below, the observed differences between the variants does not largely depend on this choice. Most notably, both the M65I and S97L mutants exhibit an altered pattern of affected residues near both mutation sites. All three variants seem to retain the cluster of residues with a large Rex contribution in the N-terminal region of the ɑ-helix, albeit with slight alterations near residue 97. The residue Leu97 of the S97L mutant shows a large R_ex_ contribution, but not Ser97 in the WT and M65I variants. Similarly, Ile65 has relatively large R_ex_ only in the M65I mutant, but nearby residues in strand β5 are differently affected in all three variants. We note that although our analysis suggests that residue Phe70 only exhibits a substantial R_ex_ contribution in the mutants, its cross-peak exhibits a large R_2_ relaxation rate also in the wild type domain, but we have excluded it from our final published analysis [8] due to resonance overlap. In the N-terminal half of the domain, residues Asp39 and Val44 have R_ex_ above 2.5 Hz only in the wild type EVH1 but not in any of the mutants. The pattern close to the N-terminus is variant-specific: Asn22 is affected only in the wild-type and the S97L mutant, whereas a cluster of residues in the region 14-16 shows differently distributed high R_ex_ values in all three variants, with variations in which residues exhibit R_ex_ above or threshold of 2.5 Hz (Figures 4C and S5).

To analyze the presence of slow timescale motions in more detail, CEST experiments were also conducted, however, these did not provide conclusive results, indicating that the differences are likely solely related to internal motions with an exchange rate faster than 200 s^-1^ [24].

### Molecular dynamics analysis supports experimental observations

Multiple rounds of molecular dynamics simulations were run on the free state of wild-type and the mutant versions of the domain. On the investigated time-scale (1 microsecond) no major differences were observable between WT and mutants. RMSF values are comparable between the different structures, except for a small deviation in the loop connecting ß1 and ß2 (Figure S6). Only terminal mobility and loop flexibility appear in the modes and the rest of the structure seems to be unaffected (also observable in the initial and final models generated from MD trajectories) (Figure S7). This is in line with the inferred secondary structures which shows that the domains seem to be stable and unaltered by the mutations in general. Some variability can be seen not in the positions containing mutations, but in their vicinity, but this does not significantly affect the assigned secondary structures. The same applies to residue contacts as well, where we see minor variations in van der Waals interactions and H-bond patterns in the mutants, but not in the exact positions of the substitutions.

### Gaussian network model analysis of the EVH1 domain identifies hinge residues with experimentally supported role in dynamics and ligand binding

We have estimated the distribution of slow internal motions in the EVH1 domain structure using the Gaussian Network Model (GNM) formalism as implemented in ProDy [31]. This simplified model takes only Cɑ positions into account, thus, this analysis is not suitable to reveal any differences in the mutants with the same overall structure, as in the present case. However, comparing the GNM results, revealing general features of the EVH1 architecture, with position-specific data obtained from NMR experiments, that are, by their nature, highly sensitive to the changes of the chemical environment of the individual residues, has the potential to reveal relevant clues on the mechanistic effects of the mutations.

The first two modes identified by GNM suggest that the “upper” and “lower” parts of the domain can move relative to each other (Figure 5A). Somewhat surprisingly, the sets of hinge residues for these modes are largely located within well-defined secondary structure elements, notably, in the beta-sheets and also near the middle of the ɑ-helix. Notably, there is a short stretch between residues 59-67 that contains hinge residues for both modes, residue Met65 itself being identified as a hinge for mode 2. Even if the GNM modes only approximately describe the actual motions of the domain, it is very likely that the M65 mutation affects a residue that is part of a segment that plays a key role in both of the first two major motional modes.This is also supported by the distribution of the residues with the largest CSP and/or changes in fitted R_ex_ values in the M65I mutant, a number of which - Thr60, Thr66, Trp77 for CSP, Phe14, Met65, and again Trp77 for R_ex_ contributions - correspond to hinge sites for mode 2 in the GNM analysis. Interestingly, several sites near mode 2 hinges have altered R_ex_ contribution in both mutants, namely, Asp39 and Val44, located in the hairpin flanked by hinge sites Tyr38 and Tyr45. For the S97L variant, the observed pattern is less in line with the GNM results, as residue Leu97, exhibiting both a large CSP and an increased R_ex_ in the mutant, is not clearly associated with a hinge region, the closest position being residue 100, a hinge in mode 1 and located almost a full turn away in the helix, facing inwards (Figure 5B).

**Figure 5.**
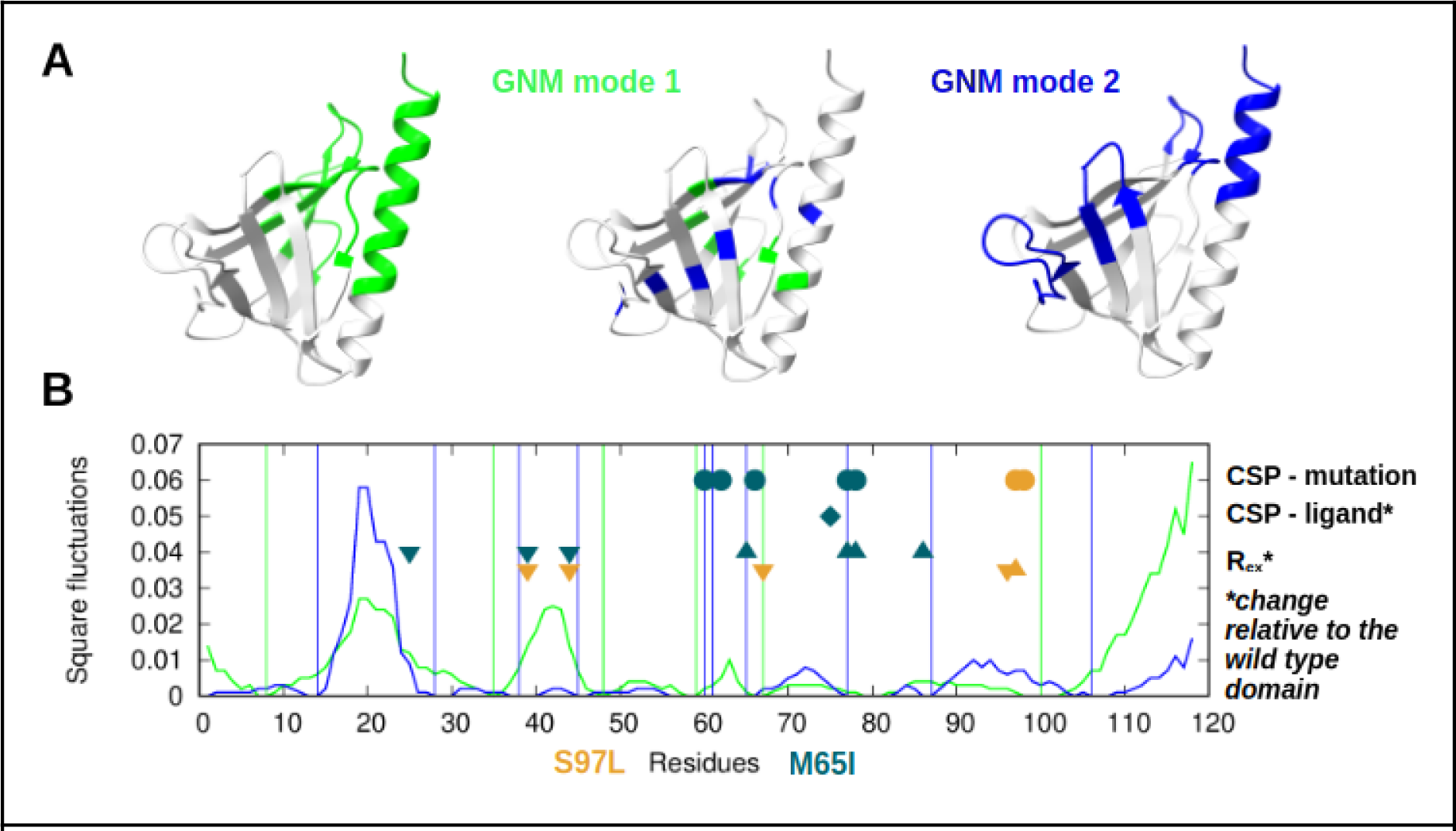
Gaussian network model analysis of the EVH1 domain architecture. **A)** In the structural models, the upper part of the structure for partitionings (left and right) according to the modes, as well as the hinge residues colored green for mode 1 and blue for mode 2. **B)** Square fluctuations along the first and second modes shown along the sequence with green and blue lines, respectively. Vertical lines denote the positions of the hinge residues of the modes. Circles show the residues with substantial chemical shift perturbation relative to the wild type domain, the diamond the perturbation upon ligand binding that differs from the wild type. Triangles indicate sites with changes in substantial R_ex_ contributions relative to the wild type, symbols pointing downwards for decrease, those pointing upwards for increase in the R_ex_. Sites are only shown where the change causes a transition to/from a large Rex (> 2,5 Hz) and the change itself exceeds 1 Hz. Symbols colored according to the mutants as shown in the key.

### Possible functional consequences of the mutations

In the InterPro database, EVH1 domains show substantial variability in the positions corresponding to the mutations M65 and S97. For M65, multiple hydrophobic amino acids occur in the various domains, but Ile is not present in this position. In the case of S97 a larger range of residues can be observed, consistent with the exposed nature of the residue (Figure S8). Interestingly, this position in Homer1 orthologues, although highly conserved, accepts Thr beside Ser, but in some species of fish Leu can also be found. These observations suggest the presence of a functional rather than structural constraint for these positions (Figure S9).

Our analysis, largely based on NMR spectroscopic data, suggests that the mutations do not alter the overall structure of the domain, rather affect its internal motions on a well-defined time scale. The nature of these motions is not exactly clear, but the distribution of the hinge residues suggests that the domain remains largely compact and the residue displacements are of moderate amplitude. The two largest motions identified by GNM divide the domain into an upper and lower parts, the lower part (in the orientation of our usual depiction) containing the ligand binding site. We speculate that the upper part might also be involved in some kind of binding events. We have noted previously the presence of a positively charged surface patch in the upper part [8], that might have some functional importance. Among the structures of related EVH1 domains, there are some examples showcasing a functional binding site in this region.

The EVH1 domain of ENAH binds one of its partners, PCARE, with a surprisingly high affinity, and the solution structure of the complex (PDB ID: 7LXF) reveals an extended binding site with key interactions far from the canonical binding site [37] (Figure 6). The ENAH P65D mutant (Pro65 in ENAH EVH1 corresponds to Thr62 in the Homer1 EVH1) showed diminished affinity, and the EVH1 of the related protein EVL, a weak binder of PCARE, could be turned to a stronger binder by introducing the Y62C and V65P mutations. These results clearly show the role of the β4-β5 loop in partner binding, and reinforce the suggestion that the M65I mutation, located also in this region, might influence additional interactions outside the canonical binding site also in Homer1 (Figure 6). Moreover, Hwang et al. note that the β5-β6 loop of ENAH EVH1 adopts a different position in the PCARE complex than in a complex with a short proline-rich peptide binding to the canonical site (PDB ID: 1EVH), and suggest that the conformation of the second binding site is key in controlling its efficiency [37] (Figure 6). This suggestion is well in line with the location of two hinge residues in the second mode identified by GNM in Homer1 EVH1, Trp77 and Gly87.

**Figure 6.**
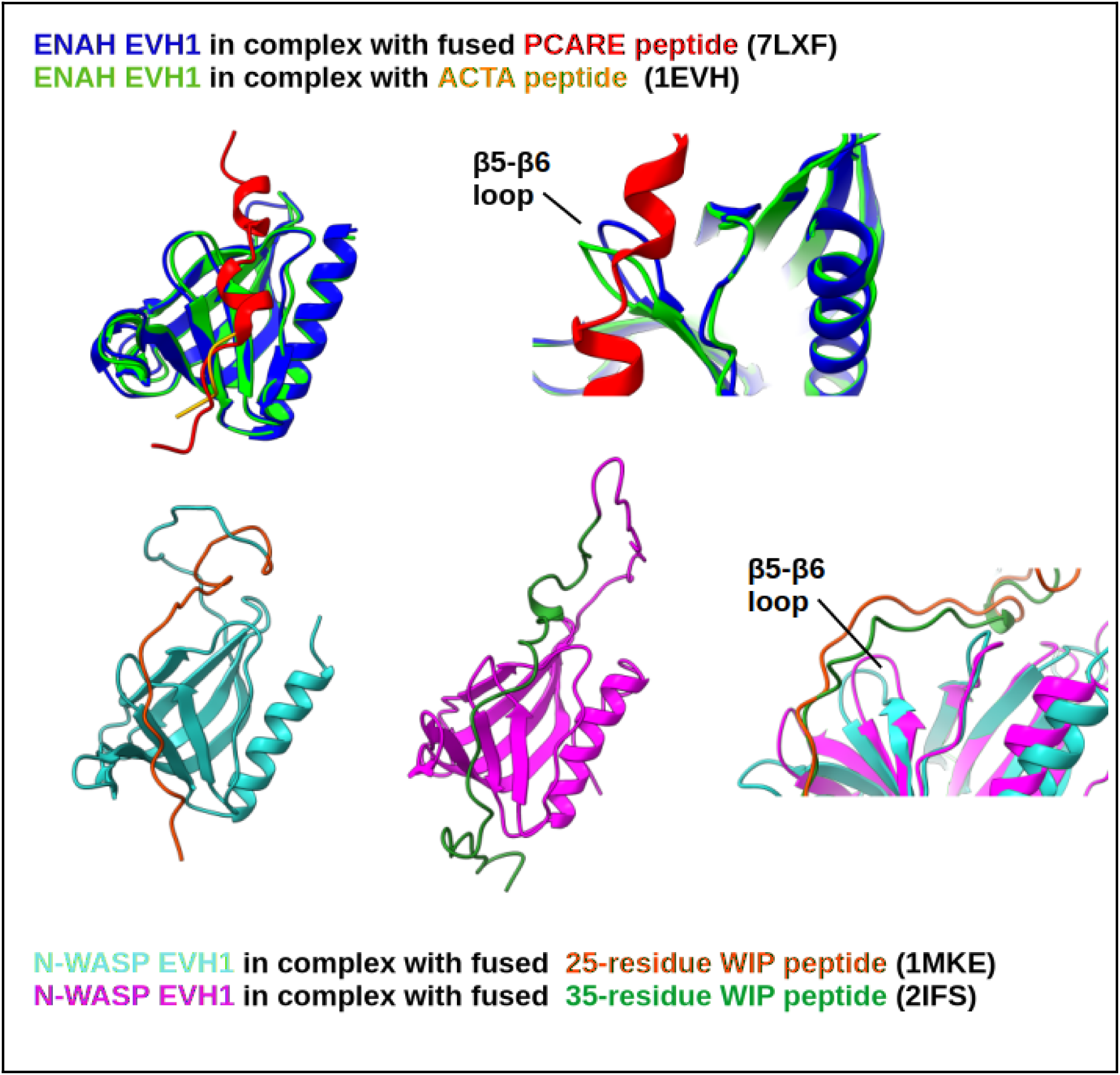
Examples of EVH1 domains with extended binding sites. The binding of longer peptide segments affects the conformation of the β5-β6 loop. See text for details. Figure prepared with ChimeraX.

In the solution NMR structures of the N-WASP EVH1 domain fused to a partner polypeptide chain derived from WIP (PDB IDs: 2IFS and 1MKE), the contribution of multiple surface regions of the EVH1 domain to the recognition is also clearly apparent. In these structures, the WIP peptide bound to the primary binding site by its Pro-rich motif wraps around the domain and makes additional contacts with the domain in the cleft between the β4-β5 and β2-β3 loops using another motif [38] [39] (Figure 6). For effective binding of WIP to the N-WASP EVH1, a much longer peptide, about 25-30 residues, is required, than typical for the Homer1 EVH1 domain. Notably, in both cases detailed above, the ENAH EVH1:PCARE and the N-WASP EVH1:WIP complexes, the binding orientation of the partner peptide is the opposite of that in known Homer1 EVH1:partner complexes (Figure 6).

Our results are compatible with a scenario where the M65I mutation exposes a cryptic binding site on the EVH1 domain. By definition, cryptic sites are not readily detectable in ligand-free structures and only appear as a consequence of internal motions [40]. Even if this is the case, the identity of the potential partner(s) remains elusive as these could only be detected by systematic screen. Another aspect is the connection between the primary and the hypothetical secondary binding sites, in other words, whether there is any kind of allosteric effect present in the domain. Third, the case of the S97L mutation is much less clear. Although this mutation also likely perturbs the internal motions, its effect is less dramatic and in several respects the opposite of that of the M65I replacement. Thus, it is by no means clear whether these two mutations, if they indeed contribute to the development of ASD as causative features, have comparable functional effects at the level of the postsynaptic protein network.

## Abbreviations

ASD: autism spectrum disorder
BLI: biolayer interferometry
BMRB: Biological Magnetic Resonance Data Bank
CD: circular dichroism (spectroscopy)
CEST: chemical exchange saturation transfer
CSP: chemical shift perturbation
EVH1: Ena/Vasp Homology domain 1
GNM: Gaussian Network Model
MD: molecular dynamics
NMR: nuclear magnetic resonance (spectroscopy)
NOE: nuclear overhauser effect
PDB: Protein Data Bank
PSD: postsynaptic density
RMSF: root mean square fluctuation
SAXS: mall-angle X-ray scattering
SDS-PAGE: sodium dodecyl sulfate-polyacrylamide gel electrophoresis
TSA: thermal shift assay

## Acknowledgements

This work was supported by the Hungarian National Research, Development and Innovation Office (NKFIH) (grants, OTKA K 137947, TKP2021-EGA-42, and 2021-4.1.2-NEMZ_KI-2022-00027). The research was also supported by the University of Debrecen, Scientific Research Bridging Fund (DETKA) and by the EU and cofinanced by the European Regional Development Fund under the projects GINOP-2.3.2-15-2016-00008 (G.B.) and GINOP-2.3.3-15-2016-00004 (access to Bruker NEO 700 MHz NMR spectrometer). We are grateful to CF Biomolecular Interactions and Crystallography of CIISB, Instruct-CZ Centre, supported by MEYS CR (LM2023042) and European Regional Development Fund-Project “Innovation of Czech Infrastructure for Integrative Structural Biology” (No. CZ.02.01.01/00/23_015/0008175). The authors thank Gitta Schlosser for the HPLC-MS measurements, and Viktor Farkas for the supervision of CD spectroscopy measurements. We acknowledge the Digital Government Development and Project Management Ltd. for awarding us access to the Komondor HPC facility based in Hungary.

## Author contributions

ZG created the concept of the study; ZG and BP designed the methodology; FF, BM, ZEK, TK, GYB, and BP performed the experiments; FF, BM, ZEK, TK, GYB, BP and ZG analysed the data; ZEK and ZG performed molecular dynamics simulations; GYB, TK and ZG provided funding and resources; ZG, FF and ZEK wrote the original draft of the manuscript; ZG, FF and ZEK created the figures and tables; and BP and GYB made manuscript revisions.

## Data accessibility

Ensembles from the molecular dynamic simulations are available at Zenodo (https://zenodo.org/uploads/17827462).

