## Supplemental Figures for "Modulation of the internal dynamics of the Homer1 EVH1 domain by putative autism-associated mutations"

for

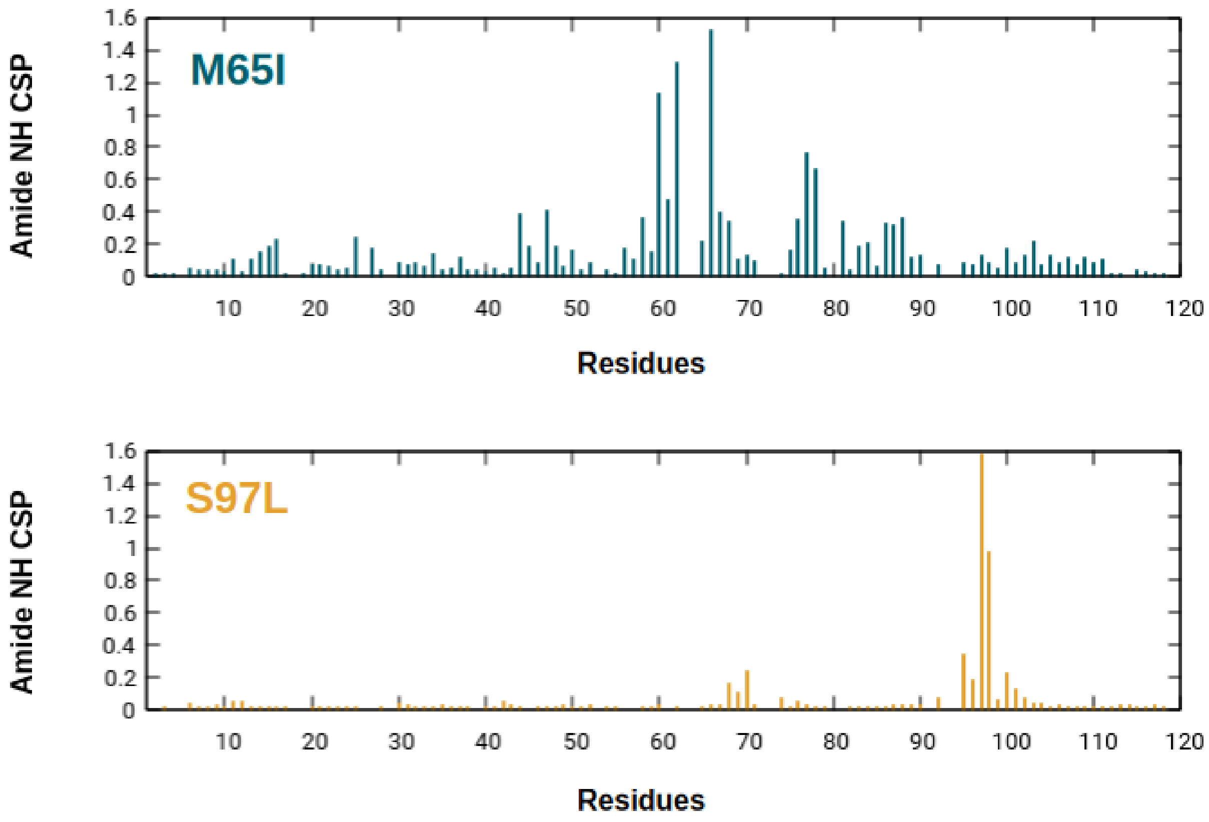


**Figure S1**. Amide NH chemical shift perturbations relative to the wild type domain (CSPs, *calculated as:* $\sqrt{0.14 * \Delta\delta N^{2} + \Delta\delta H^{2}})$ depicted separately for the two mutated domain variants investigated.

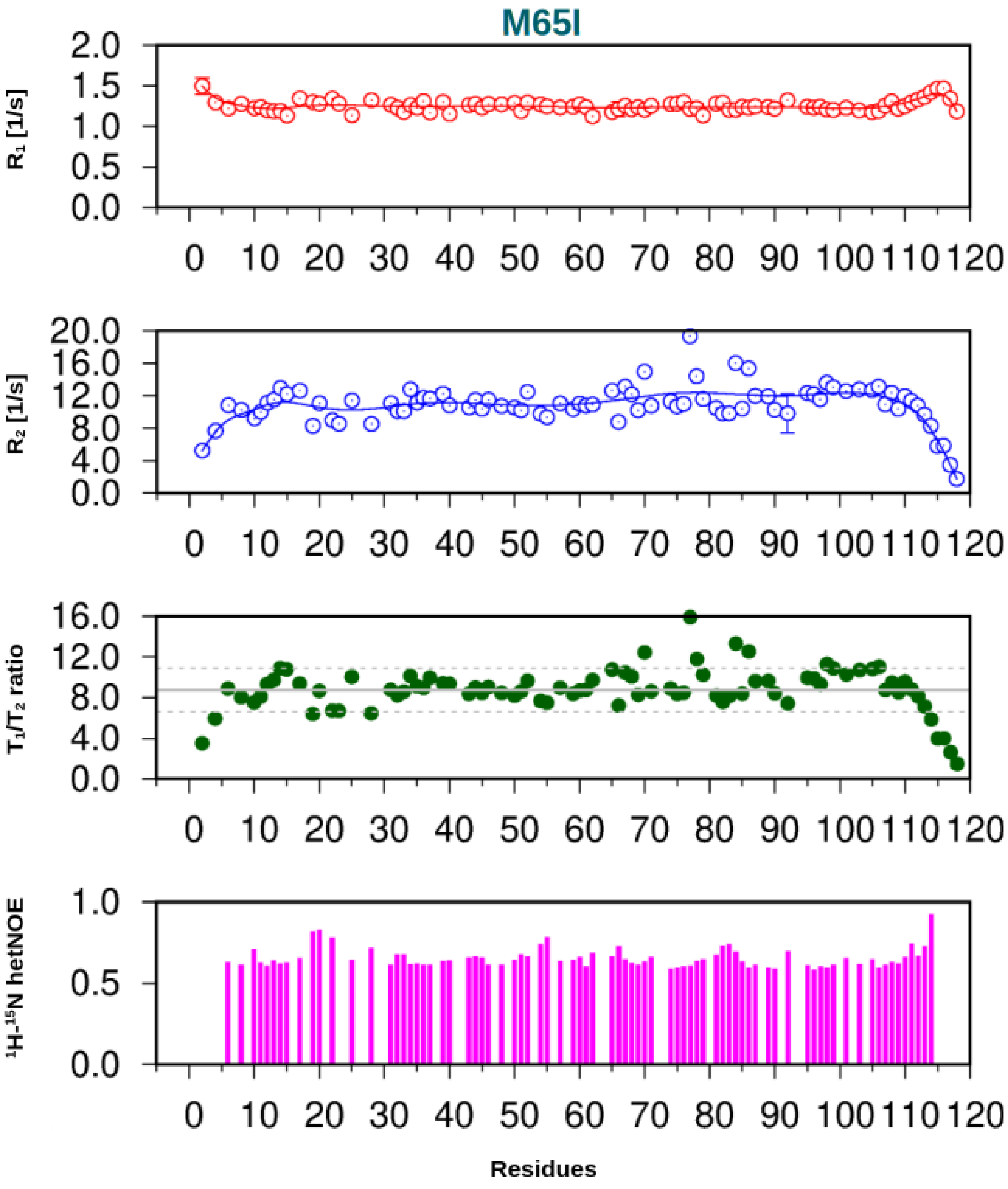


**Figure S2.** $R_1$, $R_2$ relaxation and hetNOE data for the M65I mutant. For the $T_1/T_2$ ratio, the continuous line shows the average, the dashed lines the range between 1 standard deviation. Values outside this region were not used for the calculation of the rotational correlation time.

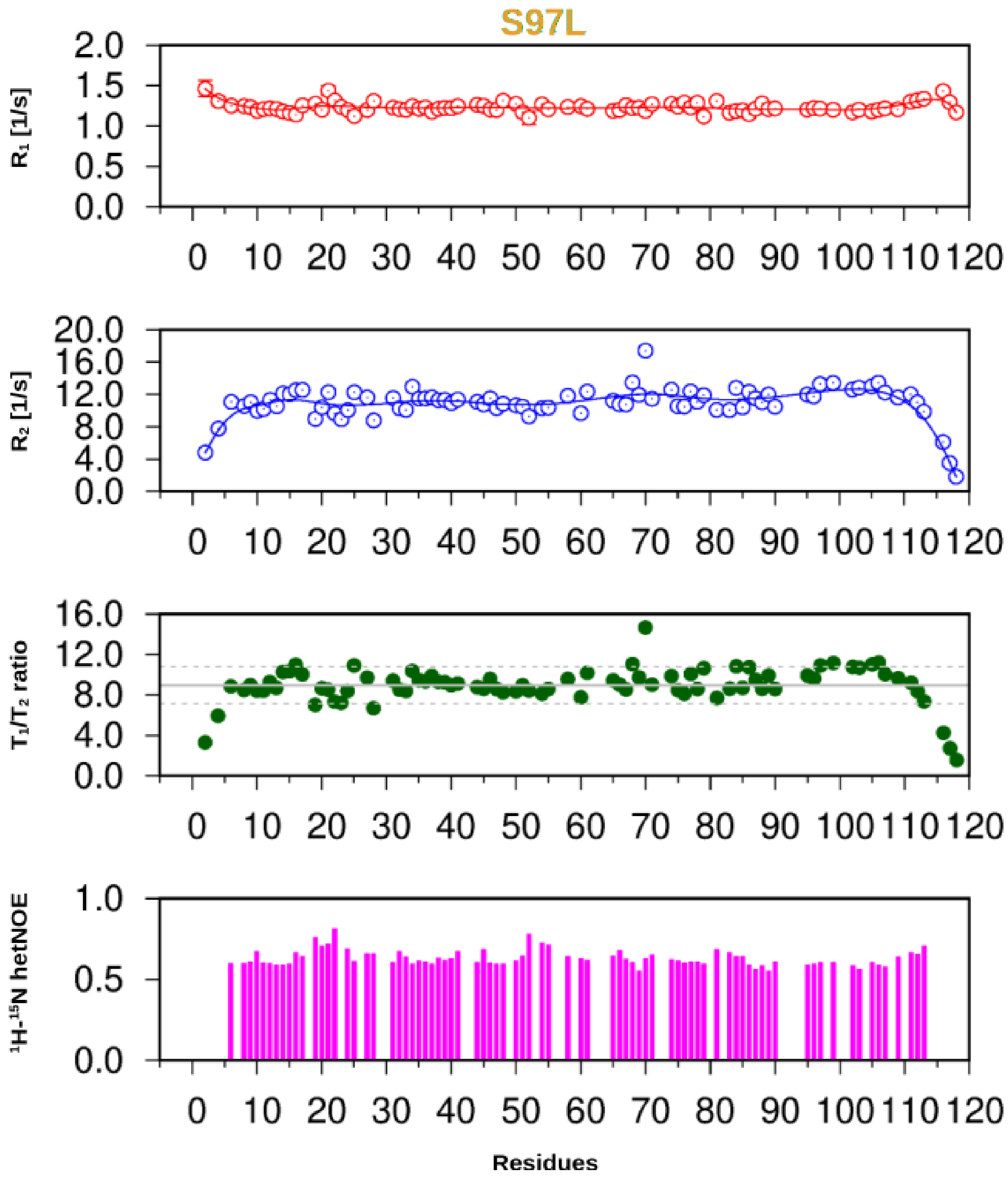


**Figure S3**. $R_1$, $R_2$ relaxation and hetNOE data for the M65I mutant. For the $T_1/T_2$ ratio, the continuous line shows the average, the dashed lines the range between 1 standard deviation. Values outside this region were not used for the calculation of the rotational correlation time.

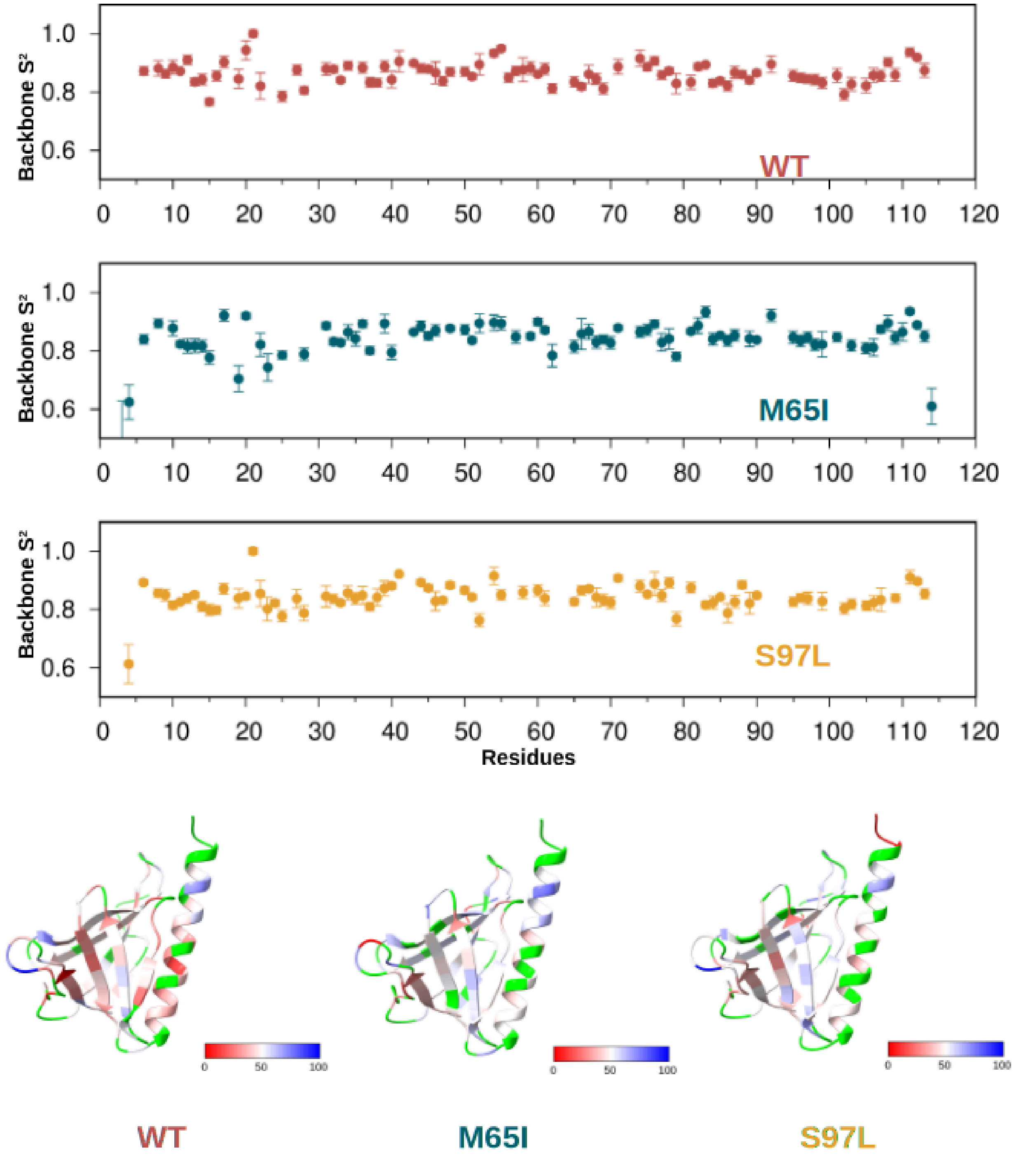


**Figure S4.** Amide $S^2$ order parameters depicted separately for the three domain variants investigated. Structural models at the bottom are colored using a red-blue scale with red corresponding to the lowest $S^2$ values within the range 0.7-1.0 (lower values are also shown in red). Residues with no fitted $S^2$ value are shown in green. Data for the wild type EVH1 domain are taken from BMRB entry 34990 (Kálmán et al., Proteins).

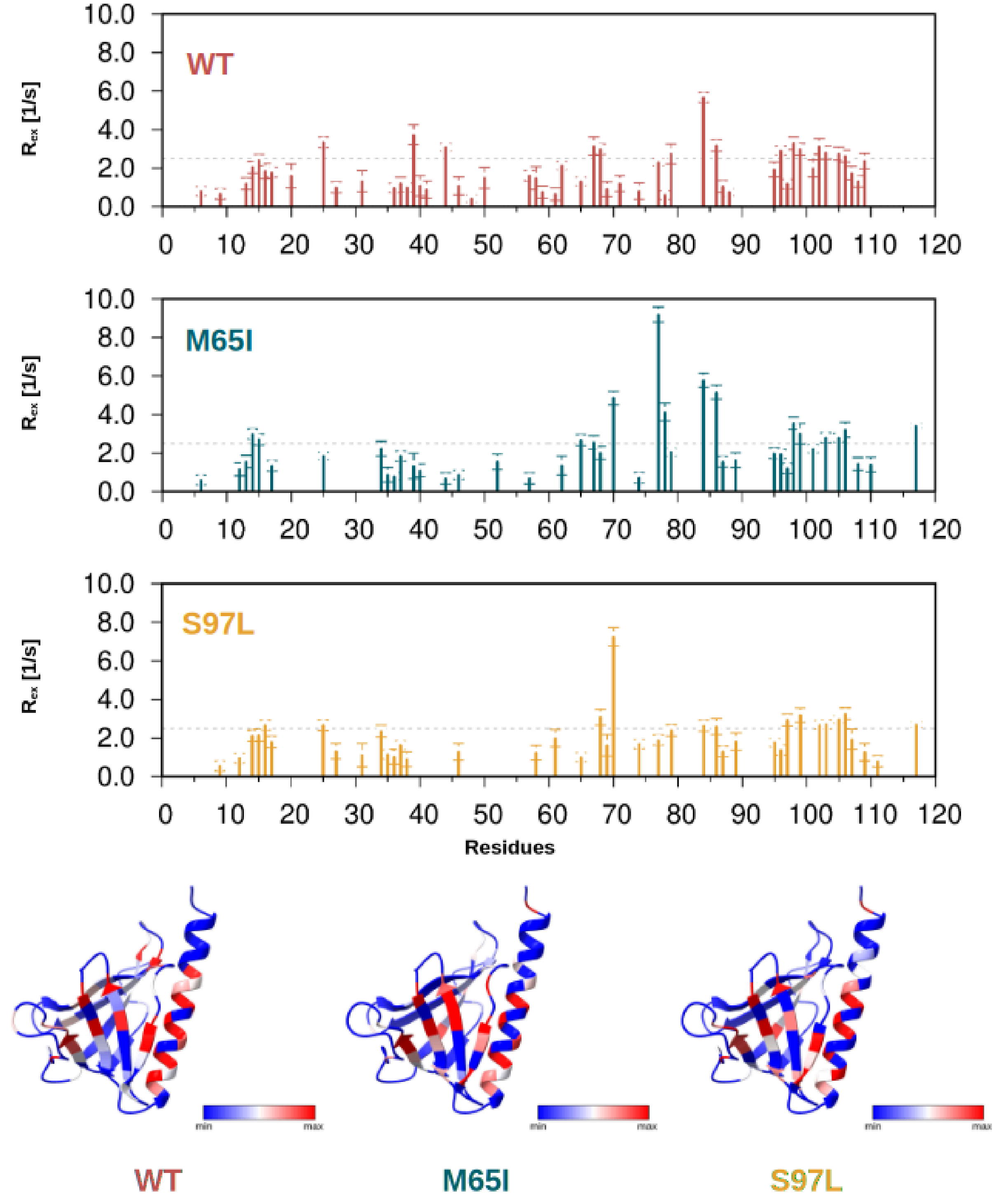


**Figure S5.** Rex contributions depicted separately for the three domain variants investigated. Structural models at the bottom are colored using a uniform blue-red scale with red corresponding to the highest $R_{ex}$ values. Data for the wild type EVH1 domain are taken from BMRB entry 34990 (Kálmán et al., Proteins).

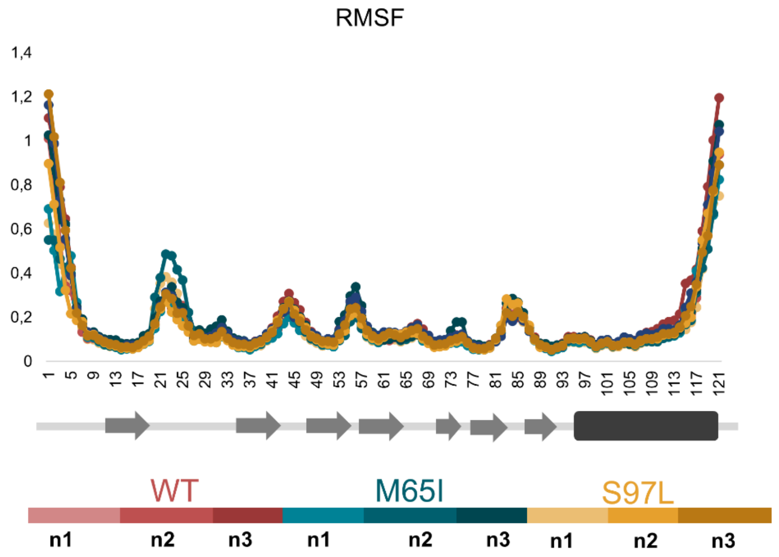


**Figure S6.** Per-residue RMSF values of the wild type and mutant structures observed during the 1 us simulations (3 runs per variant). Secondary structure elements below the chart.

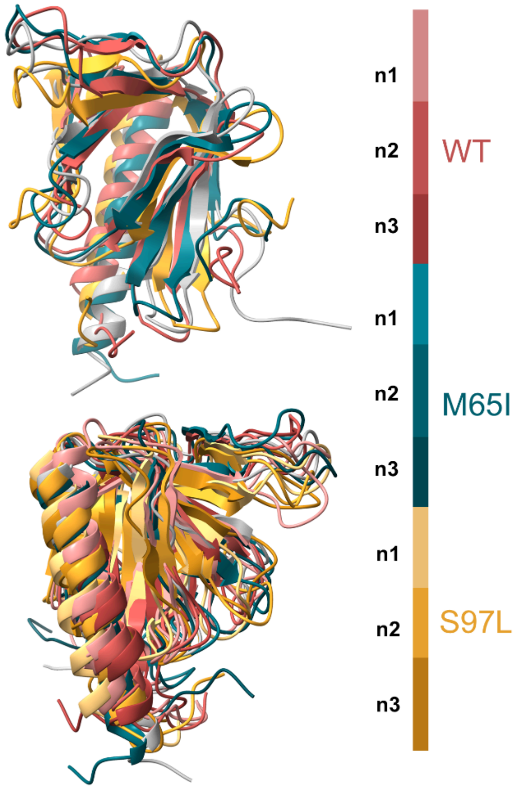


**Figure S7**. Ensembl of the models from WT and mutants. Grey represents model 1 (initial models), while the others are the last models converted from the 1 microsecond simulations. WT and mutated structures are colored according to the scheme on the right.

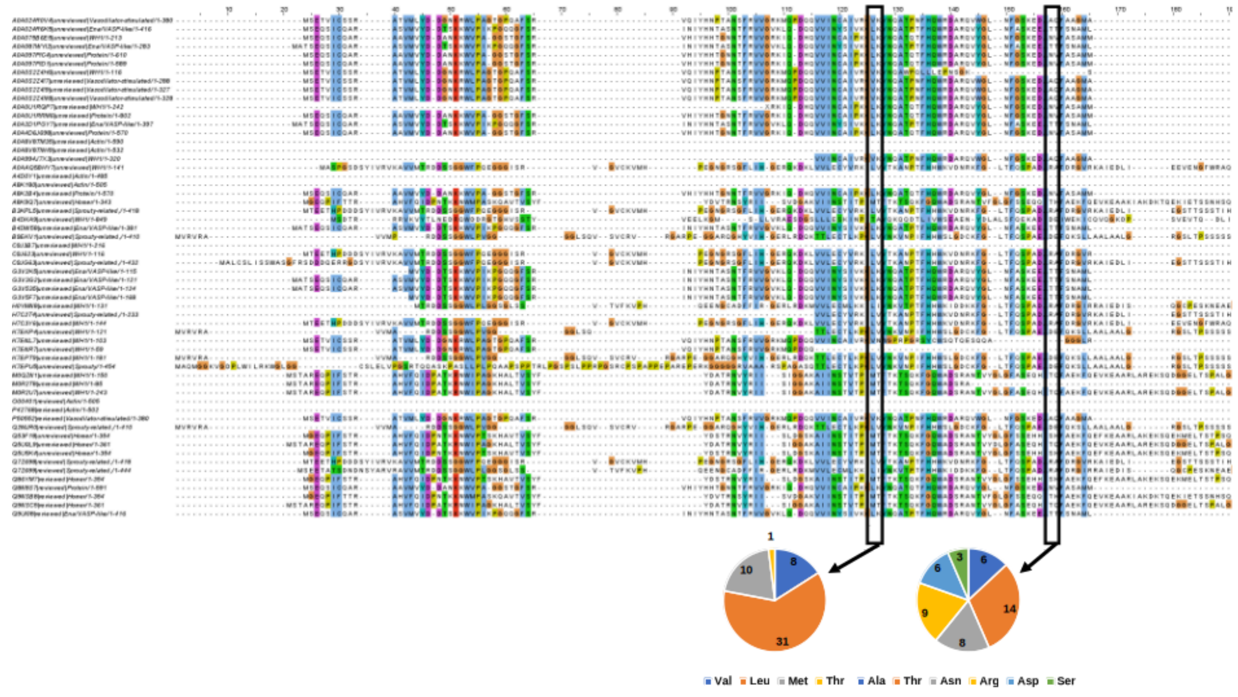


**Figure S8**. Part of the multiple sequence alignment of the human EVH1 domains in InterPro. The positions of the two mutations are highlighted and the distribution of the amino acid residues in those positions is shown on the diagrams below.

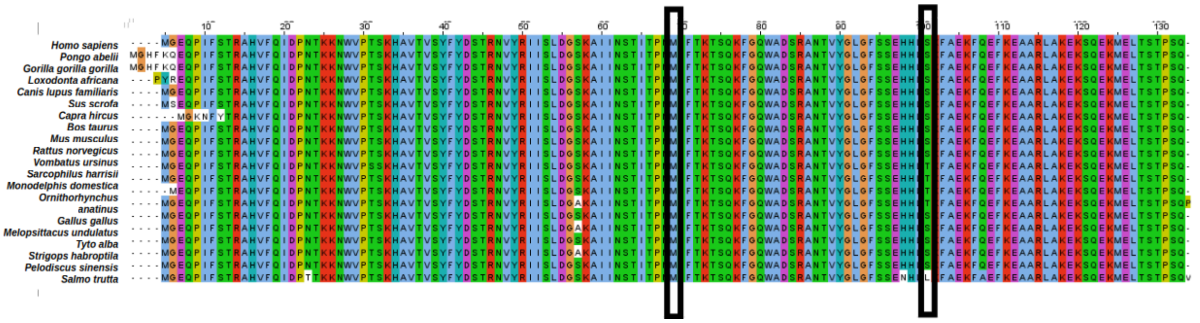


**Figure S9**. Alignment of Homer1 orthologs from different species with the positions of the mutated sites M65 and S97 highlighted.
